# New Radiological Classification of Glioma and validation with the survival analysis

**DOI:** 10.1101/2020.02.28.969493

**Authors:** Akshaykumar Nana Kamble, Nidhi K Agrawal, Surabhi Koundal, Salil Bhargava, Abhaykumar Nana Kamble

## Abstract

Radiology based classification of glioma independent of histological or genetic markers predicting survival of patients is an unmet need. Until now radiology is chasing these markers rather than focussing directly on the clinical outcome. Our study is first of its kind to come up with the independent new radiological classification of gliomas encompassing both low-and high-grade gliomas under single classification system.

TCGA-LGG and REMBRANDT public domain dataset of glioma were analyzed as training and testing dataset respectively. Based on MRI images, gliomas were classified into six types in detailed classification & three types in simplified classification system. Survival analysis using Kaplan Meier and Cox regression was done. Secondary objective was to evaluate the sensitivity and specificity of novel signs with existing histological and genetic markers.

The study predicted survival in both training and testing dataset independent of genetic or histological information. Novel signs, “Ball on Christmas tree” sign(highly specific), Type-4 lineage sign(highly sensitive) identifies IDH-wild and high-grade gliomas (grade-III and IV) while Type-2 lineage sign showed good specificity in identifying 1p19q non co-deleted IDH-mutated, ATRX del/mutated, Grade-II gliomas. There is a substantial interobserver agreement for the classification and novel signs. New radiological classification of glioma predicts the survival of patients independent of genetic or histological information. This can act as a scaffolding to formulate and streamline the treatment guidelines for glioma patients. This classification has potential of improving the quality of care of glioma patients by predicting the survival without the need of invasive biopsy.

## Introduction

It has been a century since attempts were made to classify the brain tumours. The earliest was made by P Bailey ^1^. Even being limited by the technology and statistical tests of his time, he made it a point to validate his classification with clinical data. ^(2)(3)^

The diffuse low-grade gliomas (LGGs) and high-grade gliomas are conventionally classified based on histology ^(4)^. In the old era one could learn about them after the autopsy ^(5)(6)^, or surgery ^(7) (8)^. Thus, it was too late to help patients with diagnosis made from histology, as there was no reliable and safe way of obtaining the sample from patient’s brain to predict the prognosis until the advent of radiology. Radiology with help of CT ^9^ and MRI revolutionized this workflow^(10) (11)^. Image guided biopsy is a good tool because brain unlike any other tissue of body is entirely protected by skull and lacks the redundancy, with no back up part to replace the lost ^(8)^.

The tumour types and grades help in knowing the aggressiveness of tumour in terms of survival and progression free survival. Mutations like IDH and 1p 19 q codeletion have changed our perception of looking at brain tumour^(12,13)^. These genetic markers were able to predict the survival of patients better than by conventional histological types. It has led to modification of brain tumour classification itself. Lately researchers have tried to match the radiological findings on MRI with that of genetic markers, leading to formation of ‘Radiomics’^(14)^. Its known that both in research and in clinical practice, radiological findings were matched with some invasive test (as a gold standard), which was histology some time ago and now it is genetic markers.

It is noteworthy that the histological and genetic classifications don’t consider the temporality of tumor evolution e.g. central necrosis, breakdown of blood brain barrier, appearance of satellite lesions. In our classification rather than treating low- and high-grade gliomas separately we have unified them into one classification system which is simpler to follow in clinical practice.

Knife on brain can end up hurting patient more than the disease itself. Some authors even questioned if it is justified to operate the low-grade glioma because it did not show any survival benefit over wait and watch ^(15) (9)^. We believe if we use patient’s overall survival as an external point of reference then we can have radiological classification of Gliomas as an independent predictor of survival. This would be more serviceable for developing countries where facilities of image guided interventions like stereotactic biopsies and genetic testing are not handy.

## Material and methods

The glioma databases from The Cancer Imaging Archive (TCIA) were analyzed. We used LGG-TCIA (Version 2: Updated 2016/01/05) as a training dataset and REMBRANDT (Version 1: Updated 2014/09/12) as a testing data set. All the images provided in database have been anonymized and made public since 2016/01/05 and 2014/09/12 respectively. We used these publicly available databases which included MRI images, histology data, genetic data and clinical outcome of patients. Since both databases are anonymized and in public domain, individual institutional IRB approval is not needed.

The statistical analysis was done by IBM SPSS 25.0. The results with p value <0.05 were considered statistically significant.

LGG-TCIA as a training dataset

Total 199 MRI examinations were available in LGG-TCGA Imaging archive. Histological and clinical information was obtained using TCIA website. (https://wiki.cancerimagingarchive.net/display/Public/TCGA-LGG#d60a6b8dd3934acca6dda6fe341289f4). Genetic information was obtained using National Cancer Institute GDC data portal (https://portal.gdc.cancer.gov/projects/TCGA-LGG) and Cancer Genome Atlas lower-grade glioma publication page (https://tcga-data.nci.nih.gov/docs/publications/lgg_2015/).

REMBRANDT as a testing dataset

Total 127 MRI examinations were available in REMBRANDT Imaging archive (https://wiki.cancerimagingarchive.net/display/Public/REMBRANDT). Histological and clinical information was obtained using TCIA and G-doc website. IDH and 1p19q mutation data is not available in REMBRANDT dataset.

The complete master-charts are provided in Training M1 and Testing M2 (supplementary appendix).

MRI images in the training dataset were analysed by the first author who formulated a classification scheme (figure 1) hypothesizing the temporal evolution of gliomas. This classification scheme classifies gliomas into 6 types. Some these types were grouped into one to have a more simplified classification containing only 3 types. This scheme is then plotted on a flowchart having easy steps to follow so that other readers who are blinded to patient’s outcome and genetic data can classify the gliomas. The inter-observer and intra-observer agreement were calculated using the kappa coefficient.

**Figure 1:**
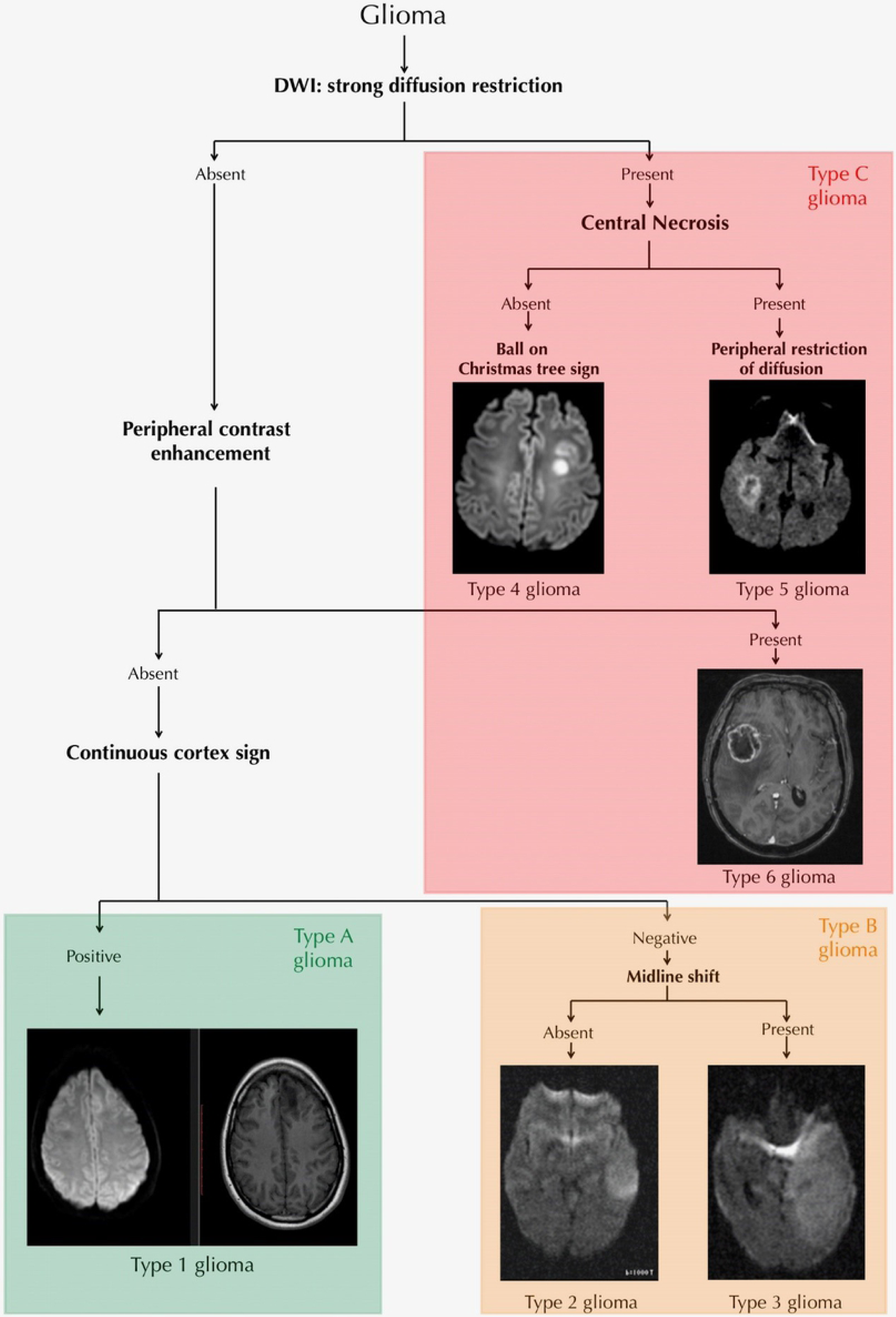
The flowchart for classifying glioma into radiological types. Simplified classification is superimposed on the detailed classification. Type A, Type B and Type C gliomas comprised of gliomas shown in green, orange and red box respectively.

The flowchart for new radiological classification of glioma is depicted in Figure 1. Steps of it are descripted below

Step 1: Whether the tumour is showing strong diffusion restriction-

If yes

Step 2A: Whether there is a central necrosis-

If No

Then it is Type 4 glioma i.e Ball on the Christmas tree sign positive.

If Yes

Then it is Type 5 glioma i.e. showing peripheral restriction of diffusion.

If tumor doesn’t show diffusion restriction

Step 2B) Whether there is peripheral contrast enhancement

If yes, then it is type 6 glioma, which is not showing peripheral restriction of diffusion but showing peripheral contrast enhancement.

If no,

Step 3) Whether there is continuous cortex sign positive

This we have accessed on T1 pre contrast images, if it’s positive then it’s Type 1 glioma.

If there is no continuous cortex sign

Step 4) Whether there is a midline shift

If no, then it is type 2 glioma

If yes, then it is type 3 glioma.

Using above mentioned flowchart gliomas were classified into 6 types, which was coined as ‘Detailed new radiological classification of glioma’.

Some of these types were clubbed into one to put forward the hypothesis that they represent same lineage of tumour. It was called ‘simplified new radiological classification of glioma’.

Type A: comprising of Type 1 glioma.

Type B: comprising of Type 2 and 3, i.e. type 2 lineage sign positive

Type C: comprising of Type 4, 5 and 6, i.e. type 4 lineage sign positive.

Nomenclature of signs

Continuous cortex sign: the neocortex is seen intact on T1 weighted images.

Ball on Christmas tree sign: The lesion is showing diffusion restriction without central necrosis seen seemingly hanging from neocortex as if from Christmas tree.

Type 2 lineage sign: It is considered as positive if glioma is classified as type B.

Type 4 lineage sign: It is considered as positive if glioma is classified as type C.

Survival analysis was done using Kaplan-Meier curve and Cox regression analysis.

Overall survival (OS) analysis was performed for both datasets. Because progression free days data was available in training dataset (LGG-TCIA), the progression free survival (PFS) analysis was also performed for this dataset.

Cox regression analysis was done by considering the possible confounding variables which may affect survival of patient. The variable which were taken into accounts were Karnofsky performance status scale, tumor location, treatment given, age, race and gender of the patient. New radiological glioma classification was included as the only classifying variable to access if novel classification can predict the survival despite of effects of confounding variables. Wald test was used to test the significance of regression coefficients of individual variables. Likelyhood ratio test was used to test significance of overall model.

Histological grading was available in both datasets thus the novel radiological signs were compared to established histological grading of glioma in both datasets. Because the genetic dataset was available in training dataset (LGG-TCIA), the novel radiological signs were compared to established genetic markers (e.g. IDH and 1 p 19 q mutations) and also with other genetic markers (e.g. ATRX and P53 mutations). This was secondary objective of our study, to compare the novel radiological signs with the established genetic markers and histological grades of glioma, to evaluate the agreement between genetic and histological classification of glioma & proposed radiological classification. The specificity and sensitivity of novel signs are calculated by cross table analysis.

## Results

New radiological classification of glioma predicted survival in both training and testing dataset as detailed in table 1. Type C of simplified classification and Type 4, 5, 6 of detailed classification had the worst overall prognosis in both datasets (log rank test, p value <0.05).

**Table 1:** showing summary of Kaplan Meier survival analysis for new radiological classification along with interobserver agreement for new radiological classification.

| Simplified new radiology classification |  |  |  | Detailed new radiology classification |  |  |  |
| --- | --- | --- | --- | --- | --- | --- | --- |
|  | Training dataset overall survival (in days) | Training dataset Progression free survival (in days) | Testing dataset overall survival |  | Training dataset overall survival (in days) | Training dataset Progression free survival (in days) | Testing dataset overall survival |
| **Type A** | 2300 | 1813 | 1653 days (55.1 months) | **Type 1** | 2300 | 1813 | 1653 days (55.1 months) |
| **Type B** | 2762 | 1460 | 1599 days (53.3 months) | **Type 2** | 1853 | 1084 | 1671 days (55.7 months) |
|  |  |  |  | **Type 3** | 3226 | 1737 | 1488 days (49.6 months) |
| **Type C** | 1378 | 638 | 750 days (25 months) | **Type 4** | 783 | 419 | 756 days (25.2 months) |
|  |  |  |  | **Type 5** | 1899 | 872 | 951 days (31.7 months) |
|  |  |  |  | **Type 6** | 464 | 464 | 684 days (22.8 months) |
| **Overall survival /progression free survival** | 2260 | 1313 | 1098 days (36.6 months) | **Overall survival /progression free survival** | 2260 | 1313 | 1098 days (36.6 months) |
| **P value** |  |  |  | **P value** |  |  |  |
| **Log Rank** | 0.004 | 0.007 | 0.022 | **Log Rank** | 0.005 | 0.007 | 0.120 |
| **Breslow** | 0.006 | 0.014 | 0.005 | **Breslow** | 0.026 | 0.033 | 0.036 |
| **Tarone-Ware** | 0.003 | 0.010 | 0.005 | **Tarone-Ware** | 0.010 | 0.020 | 0.040 |

| **Kappa** | Training set | Testing set | **Kappa** | Training set | Testing set |
| --- | --- | --- | --- | --- | --- |
| **Intra-observer (reader 1)** | 0.910 (0.035) | 0.914 (0.042) | **Intra-observer (Reader 1)** | 0.853 (0.038) | 0.823 (0.047) |
| **Inter-observer (Reader 2)** | 0.814 (0.048) | 0.745 (0.062) | **Inter-observer (Reader 2)** | 0.629 (0.053) | 0.662 (0.057) |
| **Inter-observer (Reader 2)** | 0.773 (0.053) | 0.891 (0.045) | **Inter-observer (Reader 2)** | 0.539 (0.057) | 0.734 (0.054) |

### Summary of Cox regression analysis

New radiological classification had significant impact on predicting the survival of patients in both datasets as detailed in table 2. Other than novel classification only Karnofsky performance scale and age of patient had statistically significant effect on survival of patients. The rest of the variables e.g. tumor location, type of treatment given (surgery, chemotherapy, radiation) to patient, sex and race had no effect on survival of patient (Wald test, Likelyhood ratio test; p test < 0.005). The survival analysis graphs of both training and testing dataset are shown in comparative fashion in figure 2.

**Table 2:**
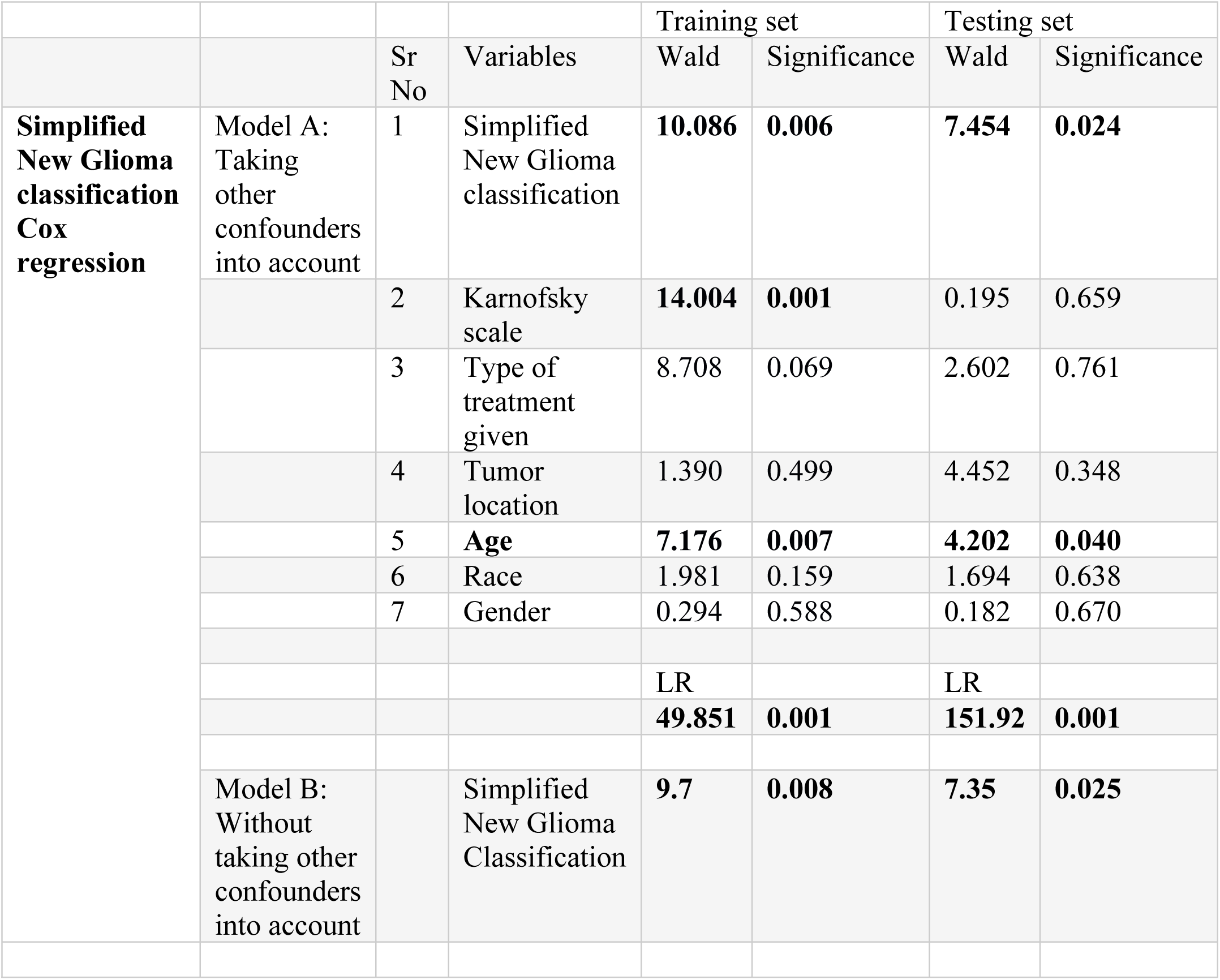
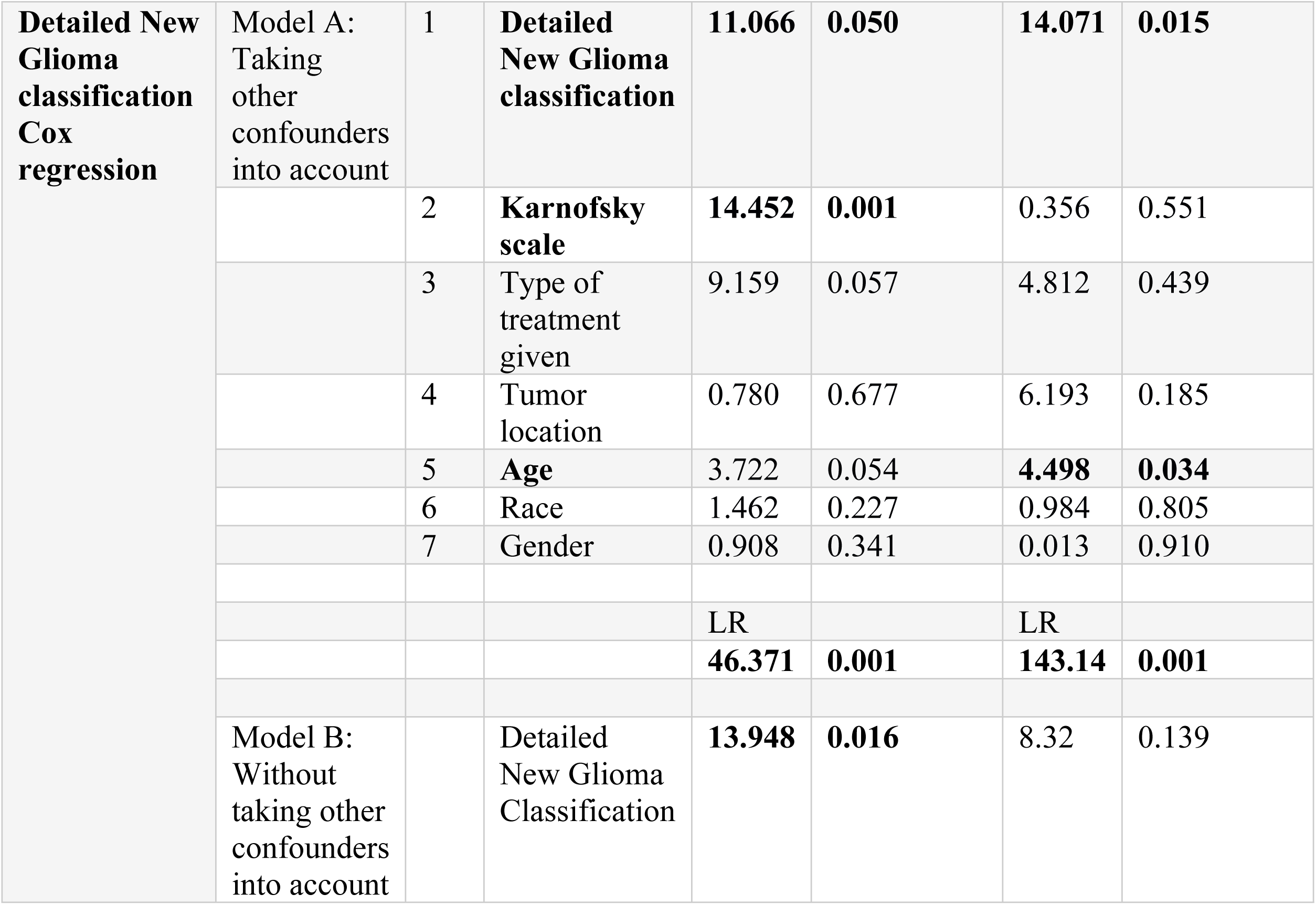
showing summary of Cox regression analysis for new radiological classification.

|  |  | Sr No | Variables | Training set |  | Testing set |  |
| --- | --- | --- | --- | --- | --- | --- | --- |
|  |  |  |  | Wald | Significance | Wald | Significance |
| **Simplified New Glioma classification Cox regression** | Model A: Taking other confounders into account | 1 | Simplified New Glioma classification | **10.086** | **0.006** | **7.454** | **0.024** |
|  |  | 2 | Karnofsky scale | **14.004** | **0.001** | 0.195 | 0.659 |
|  |  | 3 | Type of treatment given | 8.708 | 0.069 | 2.602 | 0.761 |
|  |  | 4 | Tumor location | 1.390 | 0.499 | 4.452 | 0.348 |
|  |  | 5 | **Age** | **7.176** | **0.007** | **4.202** | **0.040** |
|  |  | 6 | Race | 1.981 | 0.159 | 1.694 | 0.638 |
|  |  | 7 | Gender | 0.294 | 0.588 | 0.182 | 0.670 |
|  |  |  |  | LR |  | LR |  |
|  |  |  |  | **49.851** | **0.001** | **151.92** | **0.001** |
|  | Model B: Without taking other confounders into account |  | Simplified New Glioma Classification | **9.7** | **0.008** | **7.35** | **0.025** |
| **Detailed New Glioma classification Cox regression** | Model A: Taking other confounders into account | 1 | **Detailed New Glioma classification** | **11.066** | **0.050** | **14.071** | **0.015** |
|  |  | 2 | **Karnofsky scale** | **14.452** | **0.001** | 0.356 | 0.551 |
|  |  | 3 | Type of treatment given | 9.159 | 0.057 | 4.812 | 0.439 |
|  |  | 4 | Tumor location | 0.780 | 0.677 | 6.193 | 0.185 |
|  |  | 5 | **Age** | 3.722 | 0.054 | **4.498** | **0.034** |
|  |  | 6 | Race | 1.462 | 0.227 | 0.984 | 0.805 |
|  |  | 7 | Gender | 0.908 | 0.341 | 0.013 | 0.910 |
|  |  |  |  | LR |  | LR |  |
|  |  |  |  | **46.371** | **0.001** | **143.14** | **0.001** |
|  | Model B: Without taking other confounders into account |  | Detailed New Glioma Classification | **13.948** | **0.016** | 8.32 | 0.139 |

**Figure 2:**
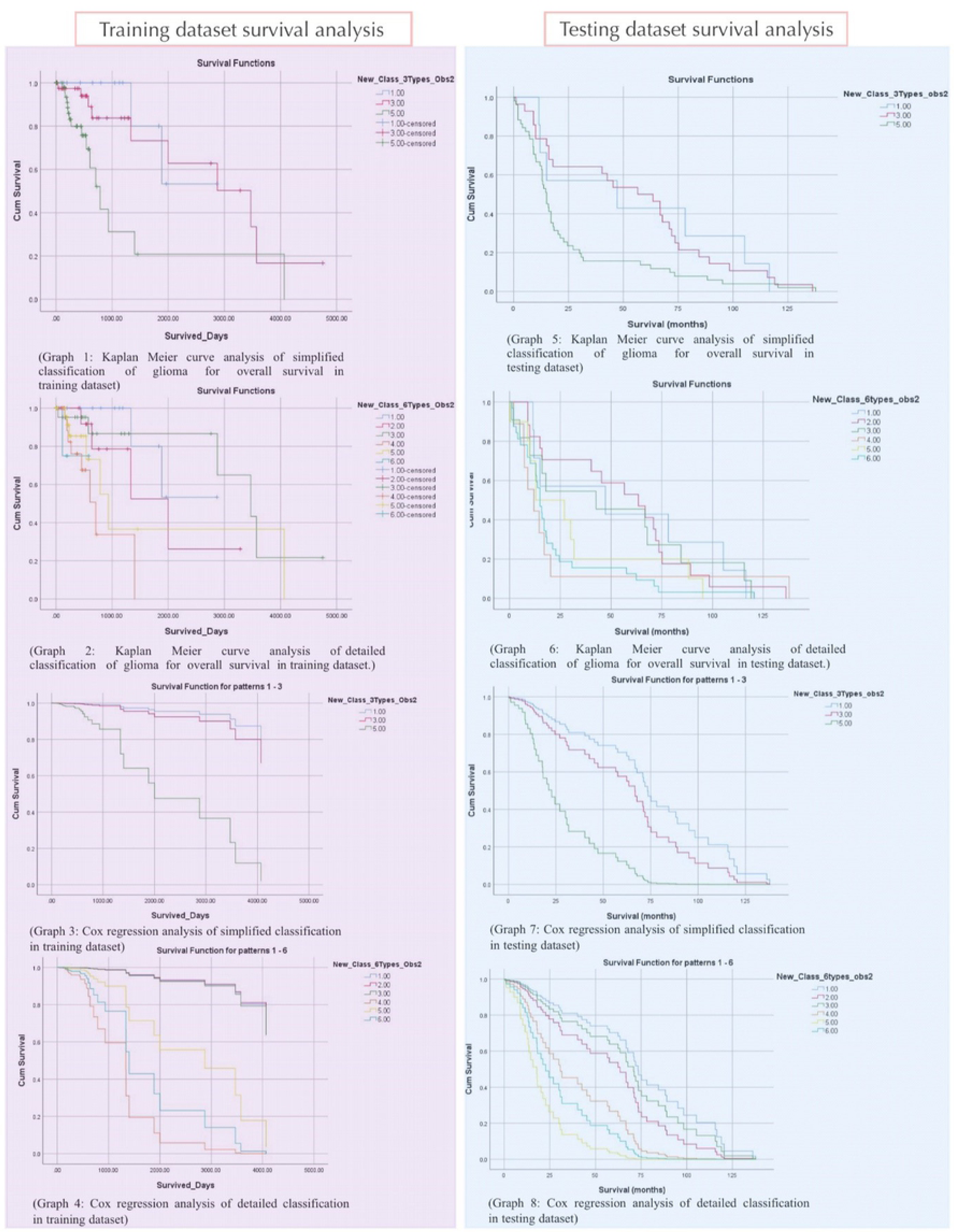
Survival analysis graphs of both training and testing dataset are shown in comparative fashion.

### Secondary objective

Novel radiological signs had a good correlation with genetic markers and histological grading of glioma as detailed in table 3. Novel signs, “Ball on the Christmas tree” sign, Type 4 lineage sign can identify IDH wild gliomas and high-grade gliomas (grade III and IV) with good specificity and sensitivity respectively (p value < 0.05); Type 2 lineage sign have good specificity in identifying 1p19q non co-deleted IDH mutated, ATRX deleted/mutated, P53 mutated and Grade II gliomas (p value < 0.05). There is moderate to substantial interobserver agreement for the new glioma classification and the novel radiological signs.

**Table 3:**
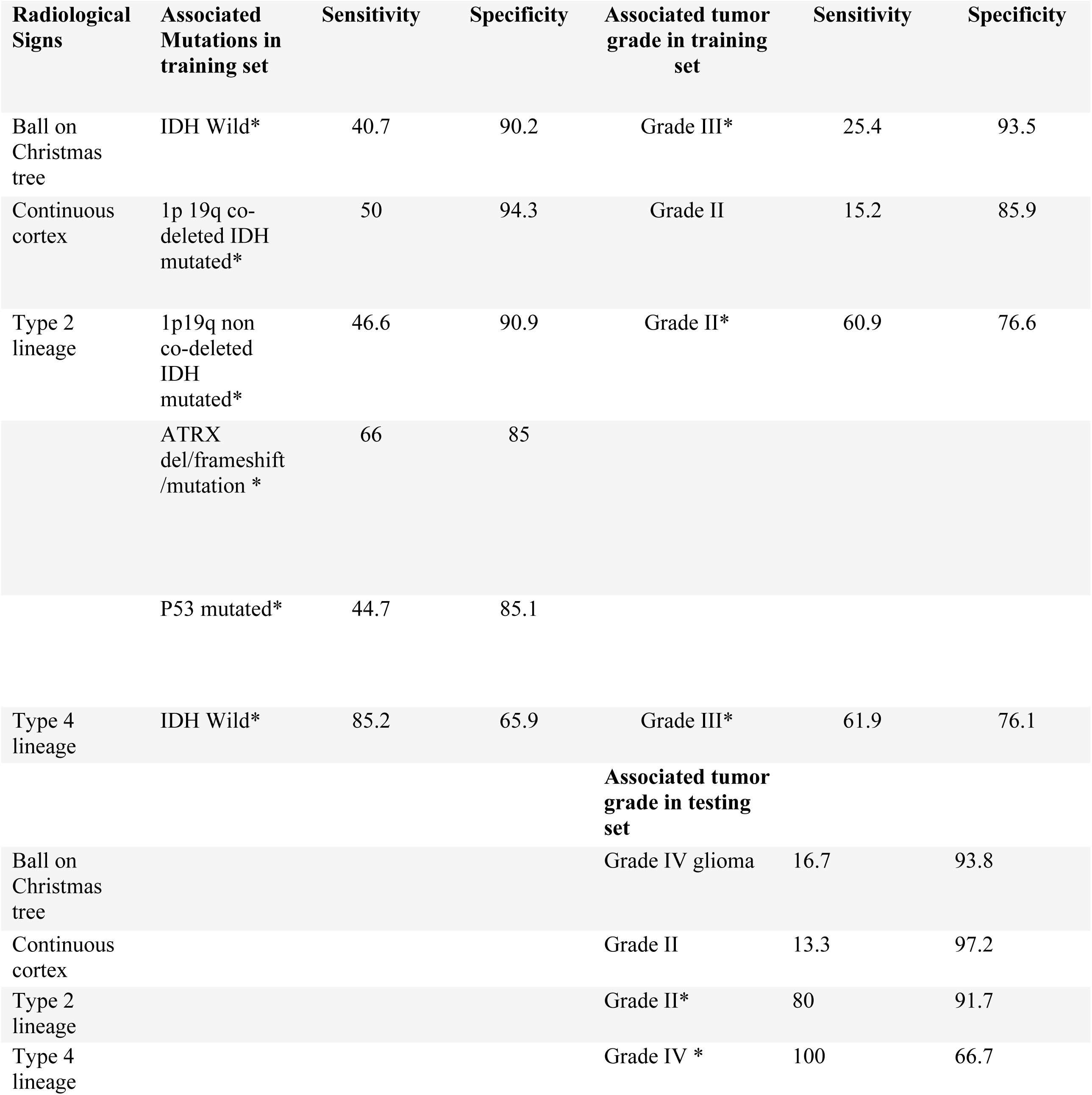
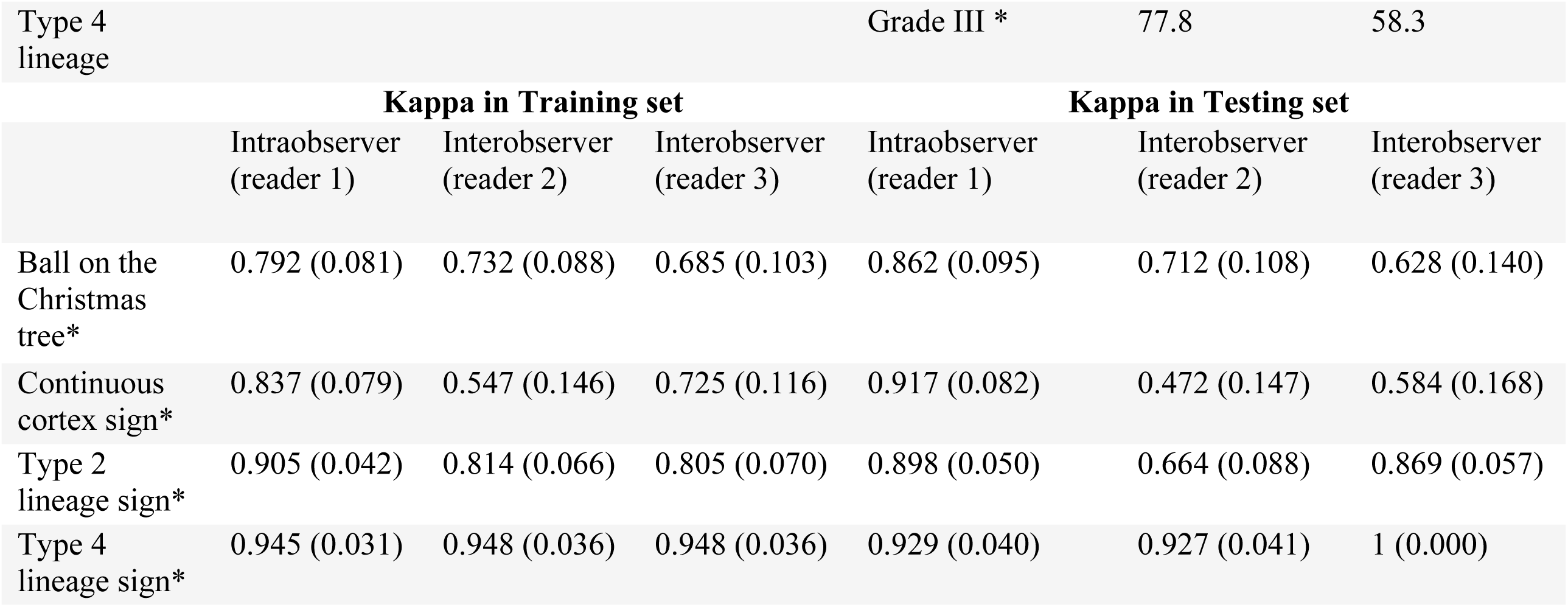
sensitivity and specificity of radiological signs in identification of genetic markers and histological grades of glioma in training dataset along with interobserver agreement for these novel signs. *p value <0.05

## Discussion

The survival analysis using Kaplan Meier curves and Cox regression analysis showed that our ‘New radiological classification of glioma’ predicted the survival of patients in both training and testing set, despite of the demographic differences in datasets. Type C gliomas based on simplified new radiology classification & Type 4, Type 5 and Type 6 gliomas based on detailed new radiology classification had the worst prognosis in both training and testing sets. The overall survival in both training and testing group are comparable for each radiological glioma type.

Cox regression analysis showed despite of presence of confounding factors the novel classification can predict the survival of patient in both datasets. Only Karnofsky performance scale and age of patient had effect on survival of patients. The rest of the variables e.g. tumor location, type of treatment given (surgery, chemotherapy, radiation) to patient, sex and race had no effect on survival of patient.

Thus, the primary objective of this study to formulate a radiological classification system for predicting survival in glioma patients independent of invasive biopsy results is achieved. The correlation between our novel radiological signs with genetic types and histological grades of glioma is also proved in our study.

Previous attempts to correlate MRI with glioma grading were based on the morphological criteria^16^ but were limited by their time because of lack of diffusion weighted images. Those studies did not give specificity and sensitivity for each grade of glioma. Even with advanced imaging, gliomas had binary division into high or low grade with no consideration for the significant survival differences between each histology grade. Despite of the differences between datasets we were able to formulate the unifying classification system and correlate with each histological grade. Our novel signs identified the Grade II, III and IV gliomas with fair sensitivity and specificity.

There have been multiple studies to correlate genetic markers with radiological signs. Patel et al described T2-FlAIR mismatch sign to identify the 1p 19q non-codeleted IDH mutant type of LGGs. It has good specificity and poor sensitivity, but little prognostic implication^17^. Many radiomics studies in radiology literature have tried to chase the elusive butterflies of the genetic markers. Our study strived to make a difference by predicting the patient’s survival. We hoped these elusive butterflies would just come and sit on our shoulders when we turn our attention to patient’s survival; thus, it was not surprising for us to find many of our novel signs had good specificity and sensitivity in detecting IDH wild gliomas, 1p 19 q non co-deleted IDH mutant, ATRX del/mutated gliomas. Reverse engineering the genotypes from the imaging phenotypes of tumours would blind us from unknown genetic markers. Our testing dataset is a good example of this, the RAMBRANDT dataset was completed before IDH mutation was in focus. Despite of having good clinical, radiological imaging data this dataset received little attention from radiology researchers after IDH and 1p 19q mutation came into spotlight. Validating with survival analysis has made our classification future-proofed, it will be relevant even with discoveries of new genetic markers.

Some studies have looked at the topography to differentiate the LGGs and association with genetic markers, e.g. frontal non-midline LGGs associated with IDH mutant LGGs ^(18)^. Some advanced imaging sequences like MR spectroscopy ^(19) (20)^ and MR perfusion^(21)^ have been used to identify high grade gliomas, but the advance imaging sequences are time consuming and some gliomas patients are too sick to lie still for longer time. Other factor is accessibility, not every institution has an access to these sequences because of expensive software packages, they need dedicated post processing to achieve standardization so that the objective results can be translated into clinical practice for decision making. Our study is not dependent on advanced sequences, we used only the basic sequences with diffusion weighted imaging (DWI) sequence to classify the gliomas. A pilot study by Wu et al have tried using DWI imaging to predict survival, but it is dependent on IDH mutation ^(22)^. Thus, none of the study have looked holistically at gliomas to come up with a radiological classification system independent of the histological and genetic information.

There is no class 1 evidence for early resection of low-grade gliomas in relatively asymptomatic patients^23^. Until now, glioma is viewed as “Shrodinger’s Cat”, requiring opening up the brain’s box to answer the patient’s survival. Glioma patient can have either good prognosis (wait and watch strategy) or bad prognosis (operation wouldn’t improve the survival). Are we justified in doing invasive biopsies in such group of patients is the key question. Our new radiological classification solves this dilemma by predicting the survival in glioma patients without the need of opening brain’s box for biopsy, thereby preventing its complications.

Some studies have used texture analysis ^(24)^ and deep learning ^(25) (26)^ to correlate the genetic marker with radiological images, but these methods can’t be applied universally. The institutions hoping to use these methods need to validate them in their own dataset, leading us back to our problem of inaccessibility to the stereotactic biopsy, genetic facilities etc. Advancements like liquid biopsies based on detecting the circulating tumor DNAs in blood are sensitive for neoplasms in other parts of body but poor for glioma ^(27)^ probably because of blood brain barrier preventing dissemination of glioma DNA. Some recent studies are using CSF to get the DNAs directly ^(28)^ but these techniques are still in research settings.

Limitations of our study includes retrospective design. Our classification is qualitative and subjective, compared to ADC (apparent diffusion coefficient) value mapping of gliomas or any other quantitative parameter. But our classification has substantial interobserver and intra-observer agreement therefore our classification can be translated into the clinical practice. Because gliomas can show necrosis (e.g. type 5 gliomas of detailed glioma classification), for ADC value mapping one needs to exclude area of necrosis which has interobserver variability in itself.

We have analysed gliomas as spectrum, ours is the first study to divide the gliomas in the lineages radiologically. This would be the first holistic classification of glioma radiologically which predicts the survival for each radiological type independent of invasive biopsy results.

## Conclusion

Proposed ‘New radiological glioma classification’ does predict the survival of patient’s with glioma independent of histological or genetic testing. This can potentially be useful in the pre-test decision making and may forgo the need of biopsy in several subset of patients. This can work as a scaffolding to formulate and streamline the treatment guidelines for glioma patients. In a tumour like glioma having poor to good prognosis, this new radiological classification can provide a valuable insight into prognosis, independent of invasive sampling procedures.

## Author contributions

Akshaykumar Nana Kamble: Conceived the idea and formulated the glioma classification. One of the radiologists who read glioma cases. Wrote the first draft. Statistics, methodology, supervision.

Nidhi K Agrawal: One of the radiologists who read glioma cases. Helped in writing the first draft. Surabhi Koundal: One of the radiologists who read glioma cases. Helped in supervision.

Salil Bhargava: Guidance and supervision. Modifying the draft.

Abhaykumar Kamble: Analysing the data. Helped in statistics. Helped in images.

## Additional information

Akshaykumar Nana Kamble: No conflict of interest.

Nidhi K Agrawal: No conflict of interest.

Surabhi Koundal: No conflict of interest.

Salil Bhargava: No conflict of interest.

Abhaykumar Kamble: No conflict of interest.

